# Integrated Artificial Intelligence and Quantum Chemistry Approach for the Rational Design of Novel Antibacterial Agents against *Ralstonia solanacearum*

**DOI:** 10.64898/2026.03.14.711561

**Authors:** Dipti Gulumbe, Goldi Tiwari, Tejas Lohar, Rohan Nikam, Amrendra Kumar, Satyabharati Giri

## Abstract

Antimicrobial resistance (AMR) in plant pathogenic bacteria poses a serious threat to global agriculture, necessitating the development of novel antibacterial agents targeting virulence mechanisms. This study presents an integrated bioinformatics-driven framework for the rational design and computational validation of Solres, a newly designed small molecule targeting key virulence proteins in phytopathogenic bacteria. Approximately 10,000 active compounds from PubChem BioAssay (AID: 588726) were analyzed using structural clustering and scaffold mining to identify conserved molecular motifs associated with antibacterial activity. Guided by high-frequency substructures, Solres was designed de novo and screened for structural novelty against PubChem, ChEMBL, and WIPO databases. Drug-likeness evaluation using Lipinski’s Rule of Five confirmed favorable physicochemical properties. Molecular docking was performed against essential virulence regulators, including PhcA, PhcR, HrpB, PehA, and Egl from *Ralstonia solanacearum* and *Xanthomonas* spp., with active sites predicted using CaspFold. Docking analyses revealed strong binding affinities and stable interactions with key catalytic and regulatory residues. Complex stability and conformational integrity were further validated through molecular dynamics simulations. Quantum chemical descriptors, including HOMO–LUMO energy gap and dipole moment, supported the electronic suitability and reactivity profile of Solres. Collectively, this study demonstrates the effective integration of cheminformatics, structural bioinformatics, molecular simulations, and quantum chemical analyses for plant-focused antibacterial discovery. The compound Solres represents a promising lead candidate for mitigating bacterial wilt disease and provides a computational framework for future experimental validation and sustainable crop protection strategies against AMR-driven phytopathogens.

## 1. Introduction

Agriculture forms the foundation of global food security, sustaining billions of people through food production, employment, and economic stability. However, modern agricultural systems are increasingly threatened by biotic stresses, particularly bacterial phytopathogens that significantly reduce crop yield and quality while disrupting soil ecological balance (Phiri et al., 2024). These pathogens not only impair plant productivity but also alter beneficial microbial communities, weaken agroecosystem resilience, and intensify reliance on chemical control strategies.

Among the most destructive bacterial pathogens, *Ralstonia solanacearum* has emerged as a globally significant threat. As the causal agent of bacterial wilt disease, it infects over 150 plant species across more than 50 botanical families, including economically vital crops such as tomato (*Solanum lycopersicum*), potato (*Solanum tuberosum*), eggplant (*Solanum melongena*), banana (*Musa* spp.), pepper (*Capsicum annuum*), and groundnut (*Arachis hypogaea*) (Wang et al., 2023; Elphinstone, 2005). Its extensive host range, environmental adaptability, and persistence make it a formidable and enduring challenge to agricultural sustainability.

*R. solanacearum* is a Gram-negative, soil-borne bacterium that enters plant roots through natural openings or wounds and subsequently colonizes xylem vessels, leading to vascular blockage, systemic wilting, and plant death (Schell, 2000). Its pathogenic success is driven by complex virulence determinants including type III secretion systems, exopolysaccharide production, quorum sensing mechanisms, and biofilm formation (Peeters et al., 2013). These systems allow the pathogen to evade plant immune responses and persist in environmental reservoirs.

The growing prevalence of antimicrobial resistance (AMR) further complicates disease management strategies. Overuse of chemical bactericides and antibiotics in agriculture contributes to the emergence of resistant strains while also disrupting beneficial soil microbiota (Ventola, 2015). Consequently, there is a critical need for innovative antibacterial agents that specifically target virulence mechanisms while minimizing environmental impact.

Recent advances in computational biology, chemoinformatics, and artificial intelligence have revolutionized drug discovery. These technologies enable rapid identification of promising molecules through virtual screening, rational compound design, and predictive modeling (Chen et al., 2018). Chemoinformatics approaches encode chemical structures using SMILES representations, allowing computational extraction of molecular descriptors and fingerprints used for quantitative structure–activity relationship (QSAR) modeling (Carracedo-Reboredo et al., 2021).

Machine learning algorithms such as Random Forest, Support Vector Machines, Logistic Regression, and Neural Networks have shown strong capability in predicting antibacterial activity based on molecular features (Liu et al., 2022). When combined with structure-based techniques like molecular docking and molecular dynamics simulations, these methods enable detailed investigation of protein–ligand interactions and stability (Durrant & McCammon, 2015).

Despite these advancements, computational agrochemical discovery remains underexplored compared to pharmaceutical research. Translating modern drug discovery frameworks into agricultural contexts offers promising opportunities for sustainable crop protection (Gadaleta et al., 2023). Computational modeling allows identification of molecular scaffolds that specifically target pathogen virulence proteins while minimizing effects on beneficial microorganisms.

In this study, we developed an integrative in silico pipeline to design and validate **Solres**, a novel SMILES-based antibacterial compound targeting virulence-associated proteins of *Ralstonia solanacearum*. By combining chemoinformatics analysis, machine learning prediction, molecular docking, molecular dynamics simulations, and quantum chemical calculations, this work demonstrates a comprehensive computational strategy for agrochemical discovery.

## 2. Materials and Methods

### 2.1 Compound Dataset Curation and Preprocessing

The primary compound dataset used for antibacterial scaffold analysis was retrieved from PubChem BioAssay (AID: 588726), deposited by The Scripps Research Institute. This dataset contains structurally annotated small molecules experimentally evaluated against bacterial targets. Approximately 10,000 compounds labeled as “active” were extracted to explore antibacterial chemical space. Each molecule was represented using its Simplified Molecular Input Line Entry System (SMILES) notation.

Structural validation and preprocessing were performed using the RDKit chemoinformatics toolkit to ensure chemical correctness and consistency. Molecules containing invalid SMILES strings, incomplete annotations, or duplicated entries were removed during preprocessing. This filtering process ensured structural integrity, minimized redundancy, and preserved chemical diversity within the dataset for subsequent scaffold analysis and compound design.

### 2.2 Fingerprint Generation and Structural Clustering

To capture molecular structural features, Extended Connectivity Fingerprints (ECFP4) were generated for all curated compounds using RDKit. These fingerprints encode topological information about atoms and their neighboring environments, enabling quantitative comparison of molecular structures.

Pairwise Tanimoto similarity coefficients were calculated between fingerprint vectors to generate a similarity matrix representing structural relationships among compounds. Based on this matrix, structural clustering was performed using the Butina clustering algorithm at a Tanimoto distance threshold of 0.4. Clusters containing at least five compounds were retained for further analysis to ensure meaningful structural representation.

Within each cluster, Maximum Common Substructures (MCS) were identified using RDKit’s FMCS algorithm. The resulting conserved structural motifs were exported in SMARTS and SMILES formats and used as templates for scaffold-guided molecular design.

### 2.3 Rational Design and Structural Optimization of Solres

A novel scaffold-inspired compound, Solres, was designed de novo based on the high-frequency MCS motifs identified during clustering analysis. The compound design incorporated structural elements associated with antibacterial activity, leading to the development of a quinoline–benzamide hybrid scaffold with balanced hydrophobic and hydrophilic properties.

The molecular structure of Solres was initially constructed in SMILES format and subsequently converted into a three-dimensional conformation using the ETKDG (Experimental Torsion Knowledge Distance Geometry) algorithm implemented in RDKit. Geometry optimization was performed using the Universal Force Field (UFF) to obtain energetically minimized conformations. The optimized structure of Solres was exported in PDB, MOL, and SDF formats for downstream computational analyses, including docking and molecular dynamics simulations.

### 2.4 Novelty and Drug-Likeness Assessment

The structural novelty of Solres was evaluated through similarity searches against publicly available chemical databases including PubChem, ChEMBL, and WIPO Patentscope to ensure that the designed compound did not significantly overlap with previously reported molecules.

Drug-likeness evaluation was performed using Lipinski’s Rule of Five, which assesses physicochemical parameters including molecular weight, hydrogen bond donors, hydrogen bond acceptors, and lipophilicity (cLogP). These parameters were used to determine whether Solres exhibits properties consistent with biologically active small molecules suitable for antibacterial development.

### 2.5 Target Selection and Protein Preparation

Major phytopathogenic bacteria were identified through literature review, including Ralstonia solanacearum, Xanthomonas spp., Pseudomonas syringae, Erwinia spp., Pectobacterium spp., and Dickeya spp. From these organisms, virulence-regulating proteins were selected as molecular targets because they play a central role in mediating bacterial pathogenicity and host infection (Genin & Denny, 2012; Mansfield et al., 2012). The selected proteins included PhcA, PhcR, HrpB, Egl, and PehA. Protein sequences were retrieved from the UniProt database, and their corresponding three-dimensional structures were obtained from the AlphaFold Protein Structure Database.

Active-site prediction was performed using CaspFold and DrugRep, which identified potential ligand-binding pockets based on structural features and residue accessibility. Prior to docking simulations, protein structures were prepared using AutoDock Tools, which involved removal of crystallographic water molecules, addition of polar hydrogen atoms, and assignment of Gasteiger partial charges. The prepared protein structures were then exported in PDBQT format for docking analysis.

### 2.6 Molecular Docking and Interaction Analysis

Molecular docking simulations between Solres and the selected virulence proteins were performed using AutoDock Vina. Docking grids were defined based on the predicted active-site coordinates identified during binding pocket analysis.

For each protein target, multiple ligand binding poses were generated and ranked according to predicted binding affinity (kcal/mol). The top-scoring complexes were selected for further interaction analysis. Visualization and structural analysis of protein–ligand complexes were conducted using PyMOL and Discovery Studio Visualizer, enabling identification of key non-covalent interactions including hydrogen bonding, π–π stacking, hydrophobic interactions, and van der Waals contacts that contribute to ligand binding stability.

### 2.7 Molecular Dynamics Simulations

To further evaluate the stability of the docked complexes, molecular dynamics (MD) simulations were performed for the highest affinity complex, particularly the PehA–Solres complex.

Simulation trajectories were analyzed using structural stability metrics including Root Mean Square Deviation (RMSD), Root Mean Square Fluctuation (RMSF), and hydrogen bond occupancy. These parameters were used to evaluate conformational stability, flexibility of amino acid residues, and persistence of ligand–protein interactions throughout the simulation period.

### 2.8 Quantum Chemical Analysis

The electronic properties of Solres were investigated using Density Functional Theory (DFT) calculations implemented in ORCA version 4.0. Geometry optimization and electronic property calculations were performed using the B3LYP exchange–correlation functional with the 6-31G(d) basis set.

Key quantum chemical descriptors were calculated, including Highest Occupied Molecular Orbital (HOMO) energy, Lowest Unoccupied Molecular Orbital (LUMO) energy, HOMO–LUMO energy gap (ΔE), dipole moment, and total electronic energy. These parameters were used to evaluate the chemical reactivity, electronic stability, and interaction potential of Solres.

### 2.9 Machine Learning Validation

To further evaluate the predicted antibacterial activity of Solres, a machine learning validation framework was implemented. A dataset comprising 20,000 compounds (10,000 active and 10,000 inactive antibacterial molecules) was compiled from NCBI, PubChem, and PubMed databases.

Multiple molecular fingerprints were generated for each compound, including FP2–FP4, DLFP, MACCS keys, ECFP2–ECFP6, and FCFP2–FCFP6. Several supervised machine learning models were trained, including K-Nearest Neighbors (KNN), Logistic Regression, LinearSVC, Support Vector Machine (SVM-RBF), Random Forest, Gradient Boosting, and Multi-Layer Perceptron (MLP).

Model training used an 80:20 train–test split, with classification accuracy used as the primary evaluation metric. To improve predictive performance, an ensemble learning framework integrating Random Forest, GLMNet, SVM-RBF, and XGBoost models was implemented.

The final predictive model was further validated using a large negative control dataset consisting of 350,000 inactive compounds, enabling robust evaluation of model discrimination and false-positive prediction rates.

### 2.10 Integrated Computational Framework

The overall computational workflow integrated chemoinformatics-driven scaffold identification, rational design of the novel compound Solres, molecular docking simulations, molecular dynamics analysis, quantum chemical characterization, and machine learning validation. This multi-layered computational pipeline provided a comprehensive in silico evaluation of Solres and facilitated identification of its potential as a novel antibacterial lead compound targeting virulence proteins of Ralstonia solanacearum.

## 3 Results

### 3.1 Novel Compound Generation

A novel antibacterial compound, Solres, was designed and computationally optimized using the RDKit chemoinformatics framework. The design strategy employed fragment-based hybridization and in silico molecular optimization to develop a quinoline–benzamide scaffold with balanced hydrophobic and hydrophilic characteristics. The resulting compound Solres, represented by the SMILES string O=C(O)c1ccc(cc1)N/N=C(\c2ccc(Nc3ccccc3)cc2)c4ccccc4.Cl, exhibited stable geometry and favorable electronic distribution, indicating potential antibacterial activity against phytopathogenic bacteria.

Structural analysis revealed the presence of a conserved C:C:C:C:C:C:N :C motif within the analyzed SMILES scaffolds, suggesting structural conservation and stability within antibacterial chemical space. The two-dimensional molecular structure of Solres, generated using RDKit, is presented in Figure 1A, while the corresponding three-dimensional conformation visualized using the MolDraw platform ([https://moldraw.com](https://moldraw.com)) (Figure 1B) illustrates the spatial arrangement, planarity, and aromatic stacking characteristics of the compound. The presence of conjugated aromatic rings and polar amide linkages within Solres suggests a strong potential for π–π interactions and hydrogen bonding with bacterial enzyme active sites. These structural and electronic features indicate that Solres represents a promising antibacterial lead compound for agricultural applications.

**Figure 1.**
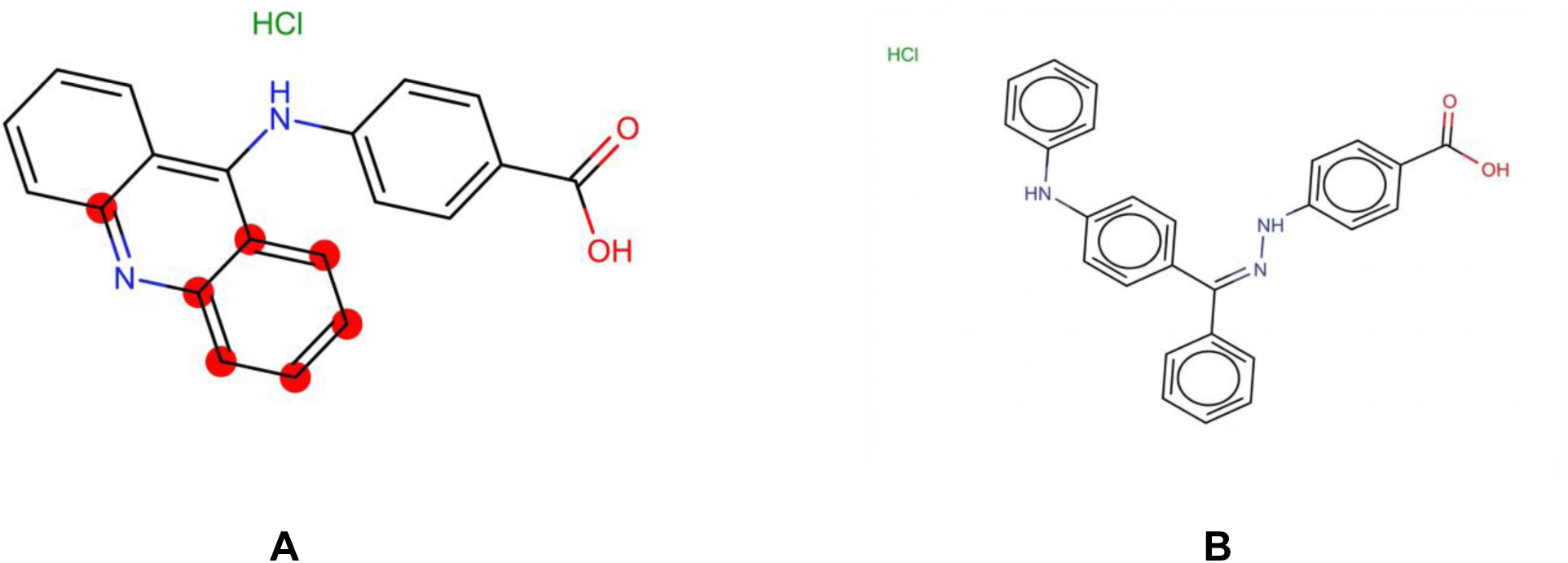
Structural representation and computational characterization of the designed novel antibacterial compound. (A) Two-dimensional chemical structure of the de novo designed quinoline–benzamide hybrid scaffold generated using the RDKit chemoinformatics framework. The structure highlights conserved aromatic regions, including the C:C:C:C:C:C:N :C motif, as well as key functional groups such as amide and carboxyl moieties contributing to structural stability and potential hydrogen bonding capacity. (B) Three-dimensional optimized conformation of the compound visualized using the MolDraw platform. The spatial arrangement demonstrates planarity of conjugated aromatic rings, balanced hydrophobic–hydrophilic distribution, and favorable geometry for π–π stacking and enzyme active-site interactions, supporting its potential as an antibacterial lead molecule.

### 3.2 Compound Validation by Lipinski’s Rule of Five

The physicochemical properties of Solres were evaluated using TargetNet ([https://nanx.app/targetnet/](https://nanx.app/targetnet/)). The compound has the molecular formula C26H22ClN3O2, with a molecular weight of 443.93 Da, 3 hydrogen bond donors, 5 hydrogen bond acceptors, and a calculated cLogP value of 7.54.

This table summarizes the molecular docking outcomes of the designed novel compound against selected virulence-associated proteins of *Ralstonia solanacearum*. For each protein target, the table presents the predicted binding affinity (kcal/mol), key interacting residues within the active site, number of hydrogen bonds formed, and major non-covalent interactions such as hydrophobic contacts and π–π stacking. The results highlight comparative binding strengths across targets, enabling identification of the most favorable protein–ligand complex for further structural and dynamic evaluation.

Assessment based on Lipinski’s Rule of Five indicated that Solres satisfies three out of the four drug-likeness criteria, including hydrogen bond donors (≤5), hydrogen bond acceptors (≤10), and molecular mass (≤500 Da). The only deviation observed was the elevated lipophilicity (cLogP > 5). Despite this violation, the compound retains significant drug-like characteristics. Increased lipophilicity may enhance membrane permeability and target binding interactions; however, it may also influence aqueous solubility. This limitation can potentially be addressed during lead optimization through structural modification, bioisosteric replacement, polar substitutions, or formulation strategies. The hydrochloride form of Solres partially mitigates solubility concerns.

### 3.3 Protein Active Site Identification

Active site prediction using DrugRep identified a well-defined central binding cavity within the AlphaFold-derived protein structures of the selected targets (Figure 2). The predicted pocket demonstrated suitable depth, solvent accessibility, and residue orientation for effective small-molecule interaction with Solres.

**Figure 2.**
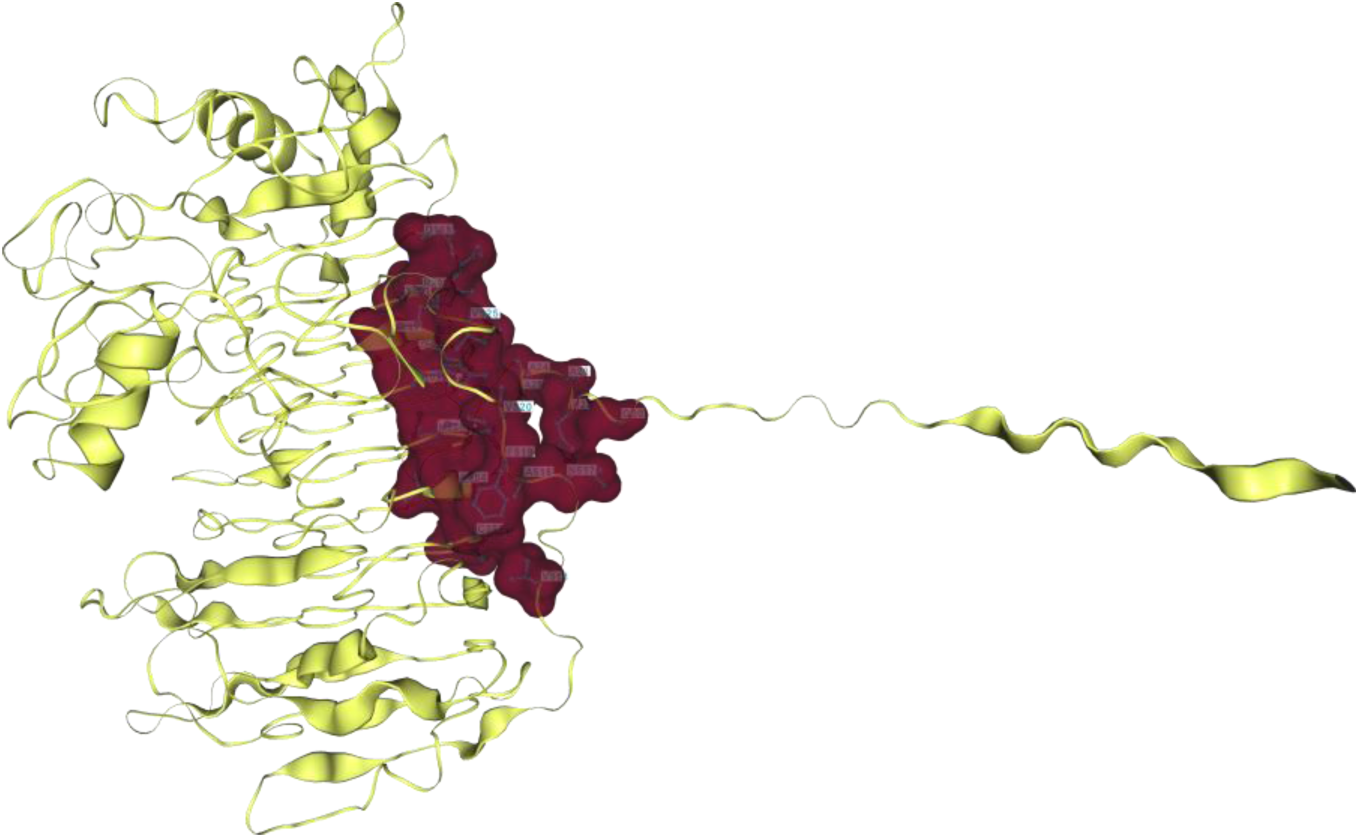
Identification and visualization of the predicted active site of the target protein.

Prior to docking analysis, protein structures were prepared using AutoDock Tools. The preprocessing steps included removal of water molecules, addition of polar hydrogens, and assignment of Gasteiger charges to ensure accurate docking conditions. The prepared protein structures were subsequently converted into PDBQT format, providing a stable and geometrically optimized framework for molecular docking simulations with Solres.

The three-dimensional structure of the target protein (**A0A2K9YUU0**) is shown in ribbon representation (yellow), with the predicted ligand-binding pocket highlighted as a surface region (red). The active site was identified using the **DrugRep Server**, which predicts potential ligand-binding cavities suitable for structure-based virtual screening. The docking grid for molecular docking was centered at coordinates **(−4.6, -16.7, -5.6)** with a grid box size of **17 × 15 × 20 Å**, covering the predicted active-site residues involved in ligand interactions. The highlighted pocket region was subsequently used to define the docking space for evaluating ligand–protein interactions and identifying potential binding conformations.

### 3.4 Molecular Docking and Visualization

Molecular docking of Solres with selected *Ralstonia solanacearum* virulence-associated proteins was performed using AutoDock (v1.1.2).

The docking analysis revealed varying binding affinities between Solres and the five target proteins, with binding energies ranging from −5.2 to −8.6 kcal/mol (Table 1). The strongest interaction was observed between Solres and the PehA protein (A0A2K9YUU0), which exhibited a binding energy of −8.6 kcal/mol. Strong interactions were also observed with PhcR (A0A0F7QWK9) and HrpB (P31778), displaying binding energies of −8.5 kcal/mol and −8.44 kcal/mol, respectively. The PhcA protein (D2WHB0) demonstrated moderate binding affinity of −7.74 kcal/mol, whereas the second isoform P52698 exhibited the lowest binding affinity of −5.2 kcal/mol.

**Table 1.**
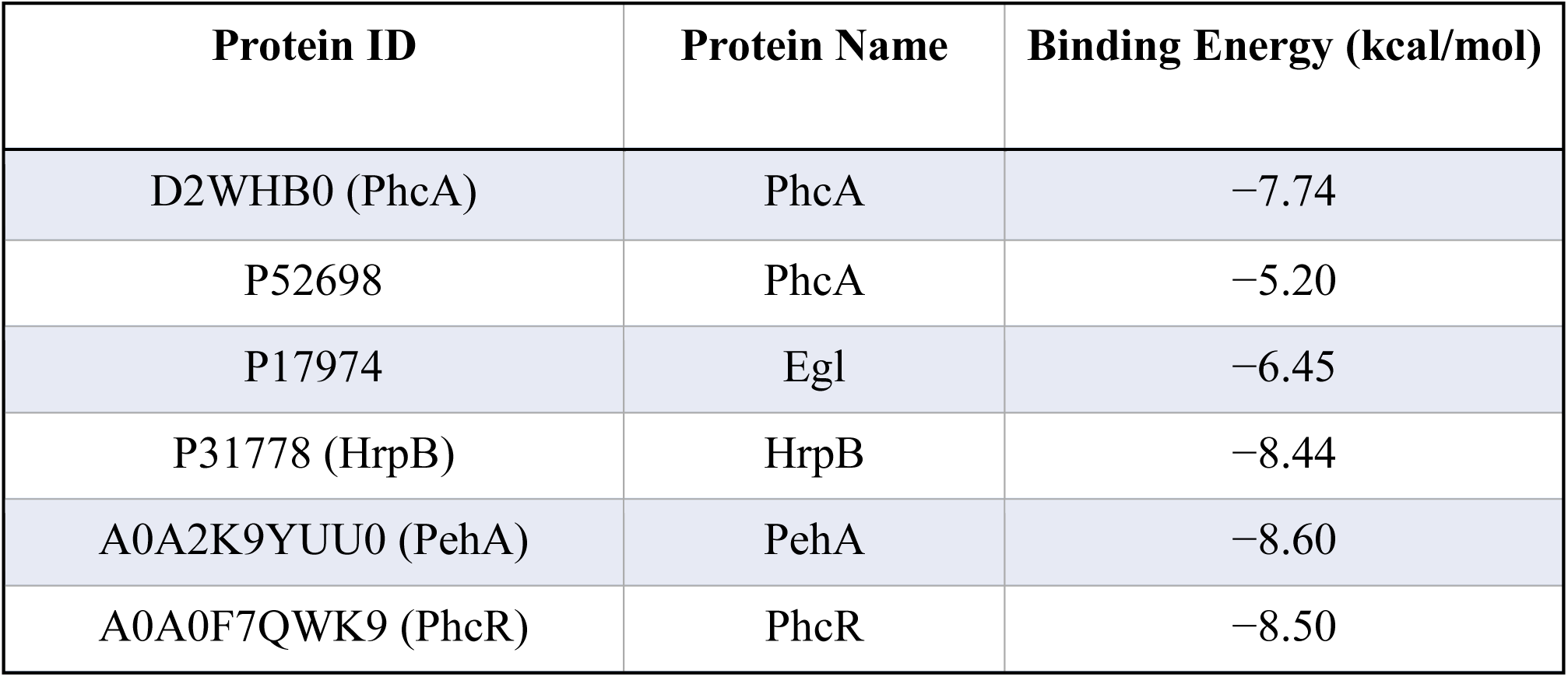
Docking results of the novel compound with selected *Ralstonia solanacearum* proteins.

These results indicate that Solres forms stable ligand–protein complexes, particularly with regulatory and virulence-associated proteins such as PehA, PhcR, and HrpB. Docking visualizations confirmed that Solres fits appropriately within the predicted active sites, forming hydrogen bonding and aromatic interactions that contribute to strong binding stability.

### 3.5 Molecular Docking Complex Visualization

The docking complex of Solres with the PehA protein was further visualized using Discovery Studio 2021 Client (version 21.1) to examine the spatial orientation and interaction profile within the active site (Figure 3A). The three-dimensional representation illustrates the PehA protein in ribbon format (brown) and Solres in stick representation (green), clearly demonstrating the accommodation of the ligand within the enzyme’s catalytic pocket.

**Figure 3.**
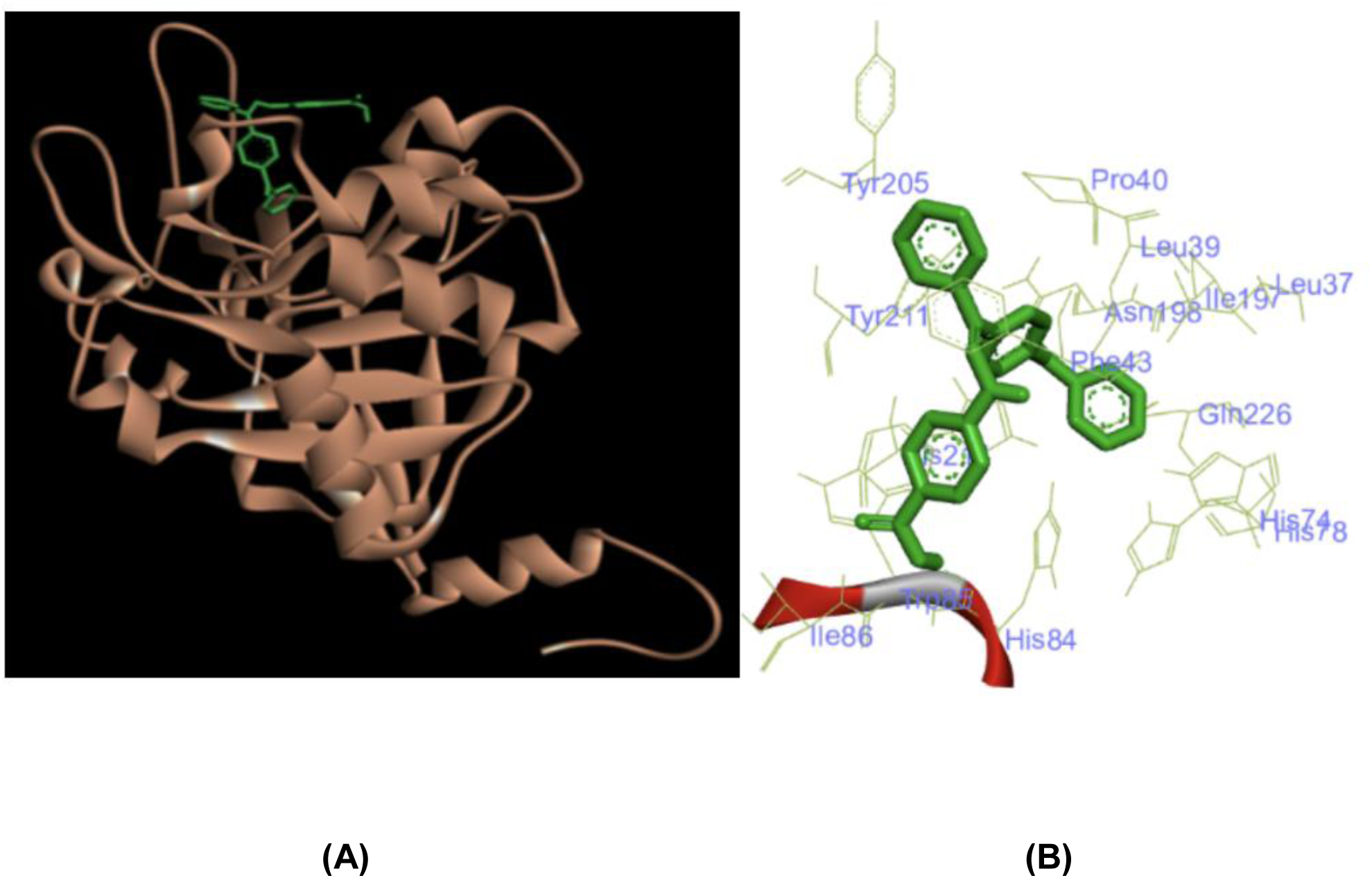
Structural visualization of the PehA–Solres docking complex. **(A)** Three-dimensional representation of the docking complex between the PehA protein of *Ralstonia solanacearum* and the designed antibacterial compound Solres, visualized using Discovery Studio 2021 Client (version 21.1). The PehA protein is shown in ribbon representation (brown) to highlight its secondary structural elements, while the ligand Solres is depicted in stick representation (green). The ligand is positioned within the predicted active-site cavity, indicating stable accommodation within the catalytic region of the enzyme. **(B)** Detailed view of the protein–ligand interaction environment showing the orientation of Solres within the PehA binding pocket. Surrounding amino acid residues including His84, Ile86, Trp85, Tyr205, Tyr211, Gln226, Pro40, Leu39, Leu37, Asn198, and Phe43 are displayed, illustrating the interaction network that stabilizes the ligand within the active site. The visualization highlights the spatial complementarity between Solres and the enzyme binding cavity, supporting the strong docking affinity observed in molecular docking analysis.

The ligand Solres is positioned firmly within the active-site cavity of PehA and interacts with the catalytic region through multiple stabilizing non-covalent interactions. The binding orientation indicates optimal spatial complementarity between Solres and the surrounding amino acid residues. This structural arrangement supports the strong docking score of −8.60 kcal/mol and suggests favorable binding stability within the enzyme’s catalytic groove.

### 3.6 Protein–Ligand Interaction Visualization

Detailed protein–ligand interaction analysis between Solres and the PehA protein of Ralstonia solanacearum was performed using Discovery Studio 2021 Client (version 21.1). Both two-dimensional and three-dimensional interaction maps were generated to evaluate the binding interactions within the predicted active-site pocket (Figure 3B).

The interaction analysis revealed that Solres forms conventional hydrogen bonds with key residues His84, Ile86, and Gln226, contributing to anchoring of the ligand within the binding cavity and enhancing complex stability. Additionally, π–π stacking and π–alkyl interactions were observed with aromatic residues Trp85, Tyr205, and Tyr211, while His214 formed a π–cation interaction, strengthening electrostatic complementarity within the binding pocket.

Further stabilization of the Solres–PehA complex was provided by van der Waals interactions involving residues Leu37, Pro40, Leu39, Asn198, and Phe43. These interactions collectively demonstrate the strong and specific binding of Solres within the PehA active site, consistent with its favorable docking score and predicted inhibitory potential.

### 3.7 Molecular Dynamics Simulation PehA–Solres Complex Stability Analysis

The structural stability of the PehA–Solres complex was further evaluated using a 100 ns molecular dynamics simulation, with conformational stability assessed through backbone Root Mean Square Deviation (RMSD) analysis (Figure 4).

**Figure 4.**
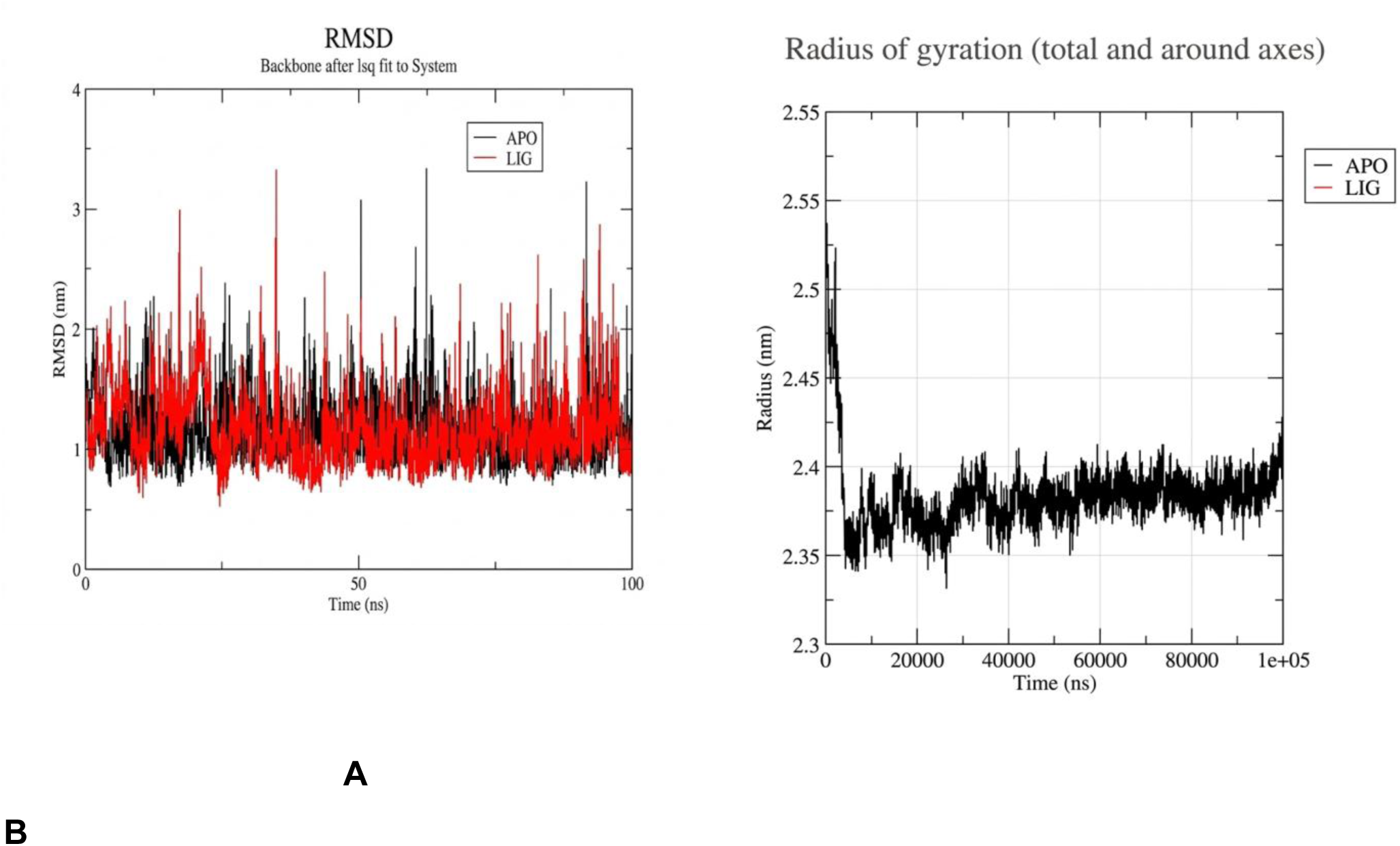
Molecular dynamics stability analysis of the PehA–ligand complex over 100 ns. (A) Root mean square deviation (RMSD) plot of the PehA protein backbone comparing the apo form (black) and ligand-bound complex (red) throughout the 100 ns simulation, illustrating structural stability and convergence over time. (B) Radius of gyration (Rg) plot of the PehA–ligand complex, demonstrating consistent structural compactness and maintenance of overall protein folding during the simulation period, indicative of stable ligand binding and conformational integrity.

Comparison between the apo protein structure and the Solres-bound complex demonstrated that both systems remained structurally stable throughout the simulation trajectory. The ligand-bound complex exhibited RMSD fluctuations within a narrow range and values comparable to or slightly lower than the apo form, indicating that binding of

The figure illustrates the spatial distribution of the highest occupied molecular orbital (HOMO) and lowest unoccupied molecular orbital (LUMO) within the protein–ligand complex. The protein structure is shown as a ribbon diagram, while the molecular orbitals are represented as colored isosurfaces (red and blue) indicating electron density distribution. The HOMO orbital is primarily localized around the ligand binding region, whereas the LUMO orbital is distributed across nearby residues of the protein structure. The energy levels associated with these orbitals are indicated, highlighting the electronic transition potential between HOMO and LUMO states.

Solres does not introduce significant structural perturbations in the protein. Importantly, RMSD values stabilized after approximately 20 ns, suggesting that the protein–ligand system reached equilibrium early during the simulation and maintained conformational stability for the remaining trajectory. These results confirm that Solres forms a stable and energetically favorable complex with the PehA protein, supporting its potential role as an inhibitor of this virulence-associated enzyme.

### 3.8 Quantum Chemical Analysis

Quantum chemical properties of Solres were analyzed using Density Functional Theory (DFT) calculations performed at the B3LYP/6-31G(d) level of theory. The calculated HOMO energy was −5.6 eV, while the LUMO energy was −2.8 eV, resulting in a HOMO–LUMO energy gap (ΔE) of 2.8 eV (Figure 5).

**Figure 5.**
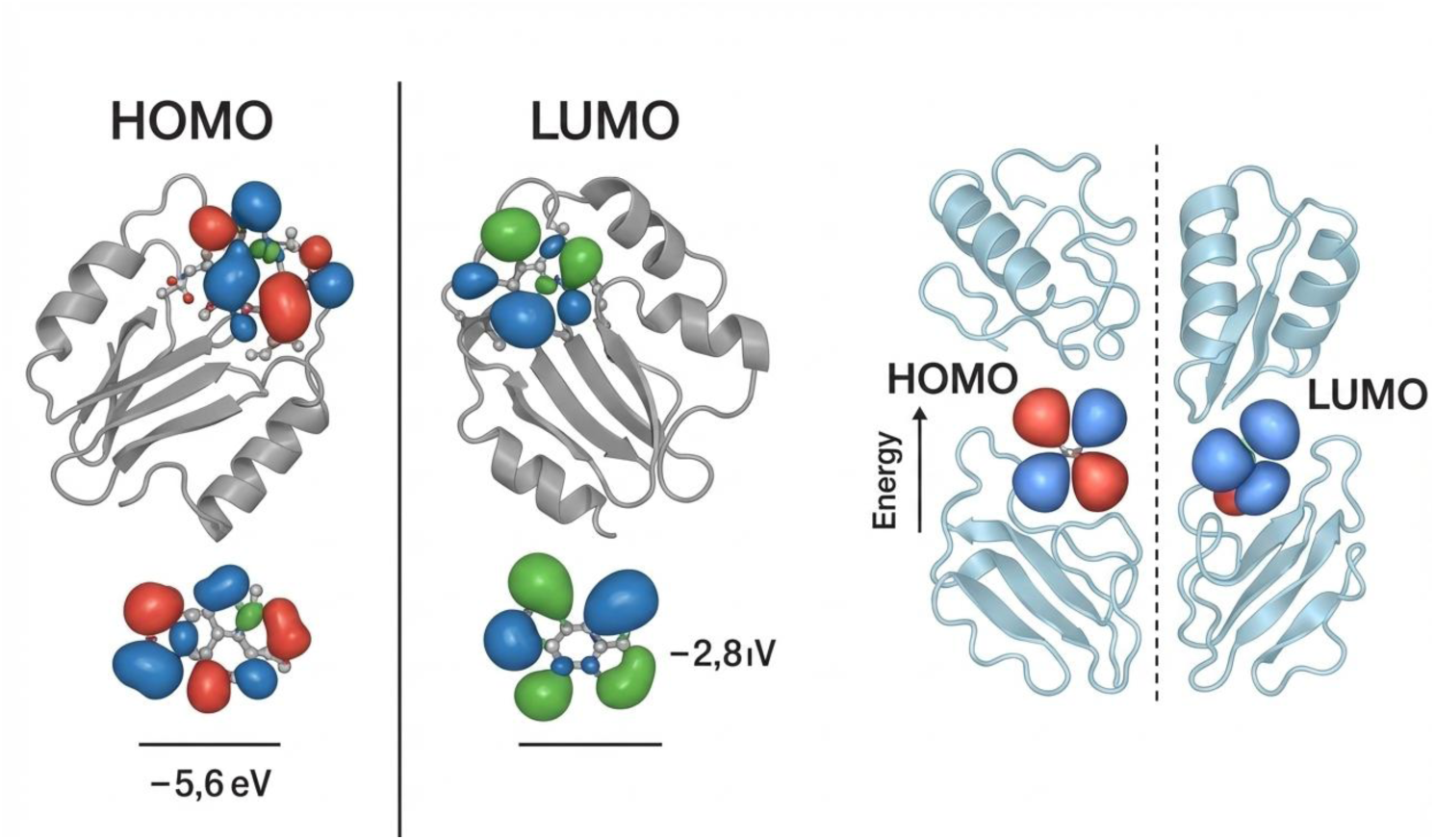
Visualization of HOMO and LUMO orbitals in the protein–ligand complex.

**Figure 6.**
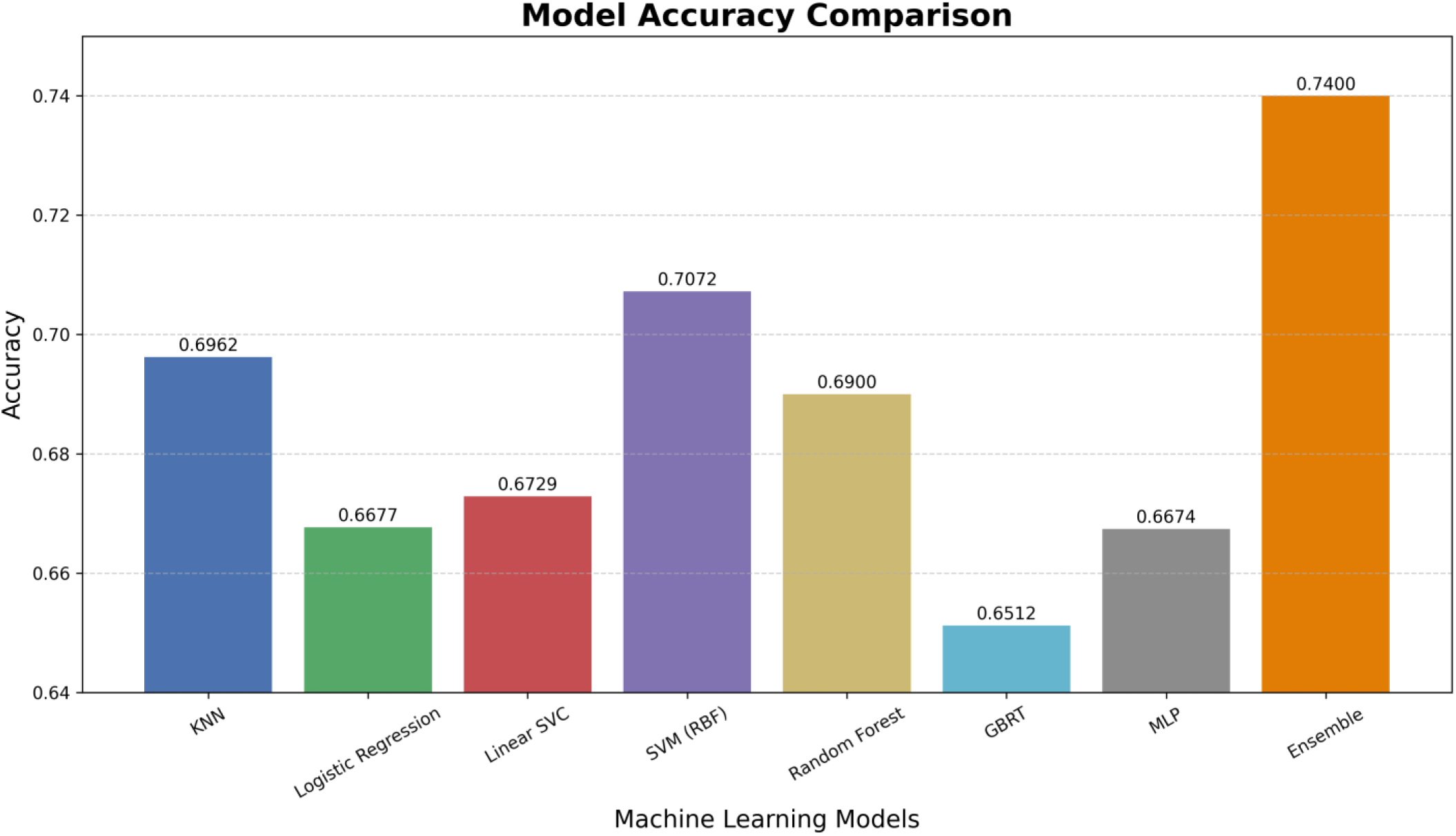
Accuracy Comparison of Machine Learning Models.

This moderate energy gap suggests a balance between molecular stability and chemical reactivity. The HOMO electron density was primarily localized over the conjugated aromatic regions of Solres, indicating strong electron-donating capacity. In contrast, the LUMO distribution was concentrated over heteroatom-containing functional groups, suggesting favorable electrophilic interaction potential. These electronic characteristics support the compound’s ability to participate in stable protein–ligand interactions.

### 3.9 Machine Learning Validation of Solres

Machine learning validation was performed to evaluate the predicted antibacterial activity

The bar chart displays accuracy scores of eight models on the dataset. The Ensemble model performs best with 0.7400 accuracy, followed by Random Forest (0.69) and SVM (RBF) (0.7072). GBRT shows the lowest accuracy (0.6512). Accuracy values are shown above each bar for clarity. The y-axis ranges from 0.64 to 0.75. of Solres using multiple molecular fingerprints, including FP2, FP3, FP4, ECFP2, ECFP4, ECFP6, FCFP2, FCFP4, FCFP6, MACCS, and DLFP.

Among the evaluated configurations, the Random Forest (RF) model using FP2 fingerprints demonstrated the highest predictive performance with an accuracy of 69%. To further improve predictive reliability, an ensemble learning model integrating Random Forest, Gradient Boosting, and Multi-Layer Perceptron classifiers was implemented.

The ensemble approach significantly improved predictive accuracy to 74%, demonstrating improved model generalization and robustness. Additional validation using a large negative control dataset containing 350,000 inactive compounds yielded an accuracy of 91%, confirming strong model discrimination between active and inactive molecules.

When Solres was evaluated using this optimized ensemble model, it was predicted as “active”, consistent with its strong docking affinity (−8.60 kcal/mol) and molecular dynamics stability. These findings collectively confirm the high likelihood of antibacterial activity of Solres against Ralstonia solanacearum, supporting its potential as a promising lead compound for further experimental validation.

## 4. Discussion

The present study demonstrates the effectiveness of integrating chemoinformatics, structure-based modeling, molecular dynamics simulations, quantum chemical analysis, and machine learning for the rational discovery of antibacterial compounds targeting phytopathogenic bacteria. Such integrative computational approaches have increasingly been applied in drug discovery to accelerate identification of bioactive compounds and reduce experimental screening costs (Lionta et al., 2014; Sliwoski et al., 2014). Using this multidisciplinary computational pipeline, we designed and validated a novel small molecule, Solres, with promising inhibitory potential against key virulence-associated proteins of *Ralstonia solanacearum*. This pathogen is responsible for bacterial wilt disease, one of the most destructive plant diseases affecting a wide range of economically important crops worldwide (Genin and Denny, 2012; Mansfield et al., 2012). The increasing prevalence of antimicrobial resistance (AMR) in plant pathogens highlights the need for innovative and targeted antibacterial strategies that minimize environmental impact while maintaining high efficacy (McManus et al., 2002; Sundin and Wang, 2018). The computational framework applied in this study provides a cost-effective and scalable approach for identifying such compounds.

The initial stage of the study involved scaffold-guided chemoinformatics analysis using a large dataset of antibacterial compounds obtained from PubChem BioAssay. Structural clustering and maximum common substructure (MCS) analysis enabled identification of conserved molecular motifs associated with antibacterial activity, an approach commonly used in ligand-based drug design (Bajorath, 2002; Vogt and Bajorath, 2012). These motifs served as the foundation for the rational design of Solres, which incorporates a quinoline–benzamide hybrid scaffold. Quinoline-based compounds have been widely reported to exhibit antimicrobial activity against a variety of bacterial pathogens (Sharma et al., 2010; Kumar et al., 2019). This structural framework combines aromatic conjugation with polar functional groups, providing a balance between hydrophobic interactions and hydrogen bonding potential. Such structural characteristics are frequently observed in biologically active antimicrobial compounds, as they facilitate both membrane permeability and strong interactions within enzyme binding pockets (Leeson and Springthorpe, 2007). The presence of multiple aromatic rings in Solres also allows for π–π stacking interactions with aromatic amino acid residues, which often play an important role in stabilizing ligand–protein complexes (Hunter and Sanders, 1990).

Evaluation of physicochemical properties using Lipinski’s Rule of Five demonstrated that Solres satisfies three of the four criteria for drug-like compounds (Lipinski et al., 2001). Although a high cLogP value may potentially reduce aqueous solubility, increased lipophilicity can enhance membrane penetration and target binding affinity (Waring, 2010). In the context of agrochemical development, moderate deviations from Lipinski’s parameters are often acceptable because agricultural applications may not require the same pharmacokinetic constraints as human therapeutics (Tice, 2001). Moreover, the lipophilicity of Solres can be addressed during lead optimization through structural modification or formulation strategies designed to improve solubility and delivery in plant systems.

Structure-based molecular docking provided critical insight into the interaction of Solres with several virulence-associated proteins of *Ralstonia solanacearum*, including PhcA, PhcR, HrpB, PehA, and Egl. Molecular docking is widely used to predict ligand binding modes and estimate binding affinities in drug discovery pipelines (Meng et al., 2011; Ferreira et al., 2015). Among these targets, the strongest binding affinity was observed with the PehA protein, which exhibited a docking score of −8.60 kcal/mol. PehA functions as a polygalacturonase enzyme involved in plant cell wall degradation during infection, making it a crucial virulence determinant for the pathogen (Huang and Allen, 2000; Tans-Kersten et al., 2001). Inhibition of this enzyme could therefore interfere with host tissue maceration and restrict bacterial colonization. The docking analysis revealed that Solres fits well within the predicted active site cavity of PehA and forms multiple stabilizing interactions with key residues. Hydrogen bonds with His84, Ile86, and Gln226 contribute to anchoring the ligand within the binding pocket, while π–π stacking interactions with aromatic residues such as Trp85, Tyr205, and Tyr211 further strengthen binding affinity. Additional stabilization through van der Waals interactions suggests that Solres achieves strong structural complementarity with the enzyme’s catalytic region.

To validate the docking predictions, molecular dynamics (MD) simulations were conducted to evaluate the stability of the Solres–PehA complex over a 100 ns trajectory. Molecular dynamics simulations are widely used to investigate protein–ligand stability and conformational flexibility beyond static docking predictions (Hollingsworth and Dror, 2018). The results demonstrated that both the apo protein and ligand-bound complex maintained stable conformational behavior throughout the simulation period. Root mean square deviation (RMSD) analysis showed that the complex reached equilibrium after approximately 20 ns and remained stable thereafter, indicating that binding of Solres does not induce significant structural perturbations in the protein. Similar stabilization effects upon ligand binding have been reported in other enzyme–inhibitor MD studies (Karplus and McCammon, 2002). These observations confirm that the interactions predicted during docking are dynamically stable under simulated physiological conditions and support the reliability of the predicted binding mode.

Quantum chemical calculations using density functional theory (DFT) further provided insight into the electronic properties of Solres. DFT methods are commonly used to study molecular reactivity and electronic properties of bioactive compounds (Parr and Yang, 1989; Jensen, 2017). The calculated HOMO–LUMO energy gap of 2.8 eV indicates a balance between chemical stability and reactivity. Molecules with moderate energy gaps often exhibit sufficient stability for biological environments while retaining the ability to participate in electron transfer interactions necessary for binding and catalytic inhibition (Geerlings et al., 2003). The distribution of the highest occupied molecular orbital (HOMO) across the conjugated aromatic regions suggests strong electron-donating capacity, while the localization of the lowest unoccupied molecular orbital (LUMO) on heteroatom-containing regions indicates potential electrophilic interaction sites. These electronic characteristics are consistent with the hydrogen bonding and electrostatic interactions observed during docking simulations.

In addition to structural validation, machine learning analysis was employed to evaluate the predicted antibacterial activity of Solres within a large chemical dataset. Machine learning has become an increasingly powerful tool in computational drug discovery for predicting biological activity and prioritizing candidate molecules (Chen et al., 2018; Vamathevan et al., 2019). Multiple molecular fingerprints were used to encode structural features, and several supervised learning algorithms were trained to distinguish active compounds from inactive ones. Among individual models, the Random Forest classifier using FP2 fingerprints achieved the highest accuracy of 69%, consistent with previous reports demonstrating the strong performance of Random Forest models in chemoinformatics classification tasks (Svetnik et al., 2003). To improve predictive reliability, an ensemble learning strategy combining Random Forest, Gradient Boosting, and Multi-Layer Perceptron models was implemented, resulting in an improved accuracy of 74%. Importantly, validation against a large negative dataset containing 350,000 inactive compounds produced an accuracy of 91%, demonstrating strong model robustness and reduced risk of false-positive predictions. When evaluated using the optimized ensemble model, Solres was predicted to be biologically active, further supporting the compound’s potential as an antibacterial agent.

Taken together, the convergence of evidence from multiple computational approaches—including chemoinformatics design, molecular docking, molecular dynamics simulation, quantum chemical analysis, and machine learning prediction—strongly supports the potential of Solres as a lead antibacterial compound targeting *Ralstonia solanacearum*. While experimental validation will be necessary to confirm biological activity and toxicity profiles, the results presented here highlight the power of integrative in silico methodologies for accelerating agrochemical discovery. By reducing reliance on traditional trial-and-error screening and enabling targeted identification of virulence inhibitors, this approach offers a promising pathway toward sustainable crop protection strategies and improved management of antimicrobial resistance in plant pathogenic bacteria.

## 5. Conclusion

This study presents an integrated computational framework for the discovery and validation of novel antibacterial compounds targeting virulence-associated proteins of Ralstonia solanacearum, a major phytopathogen responsible for bacterial wilt disease in numerous economically important crops. Using a combination of chemoinformatics, structure-based modeling, molecular dynamics simulations, quantum chemical analysis, and machine learning approaches, we successfully designed and evaluated a novel compound, Solres, as a potential antibacterial lead molecule.

Scaffold-guided chemoinformatics analysis of antibacterial compounds enabled the rational design of Solres, a quinoline–benzamide hybrid scaffold exhibiting favorable structural and electronic properties. Physicochemical evaluation indicated that the compound satisfies three of the four Lipinski’s Rule of Five criteria, suggesting acceptable drug-like characteristics with manageable lipophilicity that can be optimized during further development. Molecular docking studies demonstrated strong binding affinities between Solres and several virulence proteins of R. solanacearum, including PhcA, PhcR, HrpB, Egl, and PehA, with the strongest interaction observed for the PehA protein (−8.60 kcal/mol). Interaction analysis revealed multiple stabilizing contacts such as hydrogen bonding, π–π stacking, and hydrophobic interactions within the active site.

Molecular dynamics simulations confirmed the structural stability of the Solres–PehA complex over a 100 ns trajectory, indicating that ligand binding maintains protein conformational integrity. Quantum chemical calculations further supported the compound’s favorable electronic properties, with a moderate HOMO–LUMO energy gap suggesting balanced stability and reactivity. Machine learning validation using ensemble modeling predicted Solres as an active antibacterial compound, reinforcing its potential biological activity.

Overall, these findings highlight Solres as a promising computationally discovered antibacterial candidate and demonstrate the effectiveness of integrative in silico approaches for accelerating sustainable agrochemical discovery against phytopathogenic bacteria.

